# Potent antibody-dependent cellular cytotoxicity of a V2-specific antibody is not sufficient for protection of macaques against SIV challenge

**DOI:** 10.1101/2023.11.14.567003

**Authors:** Michael W. Grunst, Hwi Min Gil, Andres G. Grandea, Brian J. Snow, Raiees Andrabi, Rebecca Nedellec, Iszac Burton, Natasha M. Clark, Sanath Kumar Janaka, Nida K. Keles, Ryan V. Moriarty, Andrea M. Weiler, Saverio Capuano, Christine M. Fennessey, Thomas C. Friedrich, Shelby L. O’Connor, David H. O’Connor, Aimee T. Broman, Brandon F. Keele, Jeffrey D. Lifson, Lars Hangartner, Dennis R. Burton, David T. Evans

**Author notes:** Corresponding author: David T. Evans, 585 Science Drive, Madison, WI 53711. These authors contributed equally to this work (Michael W. Grunst, Hwi Min Gil).

## Abstract

Fc-mediated antibody effector functions, such as antibody-dependent cellular cytotoxicity (ADCC), can contribute to the containment HIV-1 replication but whether such activities are sufficient for protection is unclear. We previously identified an antibody to the variable 2 (V2) apex of the HIV-1 Env trimer (PGT145) that potently directs the lysis of SIV-infected cells by NK cells but poorly neutralizes SIV infectivity. To determine if ADCC is sufficient for protection, separate groups of six rhesus macaques were treated with PGT145 or a control antibody (DEN3) by intravenous infusion followed five days later by intrarectal challenge with SIV_mac_239. Despite high concentrations of PGT145 and potent ADCC activity in plasma on the day of challenge, all animals became infected and viral loads did not differ between the PGT145- and DEN3-treated animals. To determine if PGT145 can protect against a neutralization-sensitive virus, two additional groups of six macaques were treated with PGT145 and DEN3 and challenged with an SIV_mac_239 variant with a single amino acid change in Env (K180S) that increases PGT145 binding and renders the virus susceptible to neutralization by this antibody. Although there was no difference in virus acquisition, peak and chronic phase viral loads were significantly lower and time to peak viremia was significantly delayed in the PGT145-treated animals compared to the DEN3-treated control animals. Env changes were also selected in the PGT145-treated animals that confer resistance to both neutralization and ADCC. These results show that ADCC is not sufficient for protection by this V2-specific antibody. However, protection may be achieved by increasing the affinity of antibody binding to Env above the threshold required for detectable viral neutralization.

**Author Summary:** Antibodies that bind to the human immunodeficiency virus (HIV-1) envelope glycoprotein (Env) on virions can neutralize viral infectivity. Antibodies may also bind to Env on the surface of virus-infected cells and recruit immune cells to eliminate the productively infected cells through a process known as antibody dependent cellular cytotoxicity (ADCC). In rare instances, certain antibodies are capable of mediating ADCC despite negligible neutralizing activity. Such antibodies are thought to have contributed to the modest protection observed in the RV144 HIV-1 vaccine trial and in some nonhuman primate studies. One antibody, PGT145, was found to cross-react with simian immunodeficiency virus (SIV) and to mediate potent ADCC against SIV-infected cells despite weak neutralization of viral infectivity. We therefore tested if the potent ADCC activity of PGT145 could protect rhesus macaques against mucosal challenge with pathogenic SIV. PGT145 did not protect against wild-type SIV_mac_239, but did protect against an SIV_mac_239 variant with a single amino acid substitution in Env (K180S) that increases antibody binding to Env and makes the virus susceptible to neutralization. Thus, while ADCC may contribute to protection against immunodeficiency viruses through the elimination of productively infected cells, the higher affinity of Env binding necessary for potent neutralization is a critical determinant of antibody-mediated protection.

## Introduction

The RV144 trial remains the only clinical vaccine trial to show a modest reduction in the rate of HIV-1 infection [1]. Antibodies capable of binding to the variable 1 and 2 (V1V2) loops of HIV-1 gp120 were identified as a correlate of protection [2] and sequence analysis of breakthrough infections later revealed V2 signatures consistent with a sieving effect of vaccine-induced immune responses [3]. While these antibodies did not neutralize circulating HIV-1 isolates, a non-significant trend towards a lower risk of HIV-1 acquisition was observed among vaccinees with higher antibody-dependent cellular cytotoxicity (ADCC) [2]. Subsequent studies also revealed the induction of Env-specific antibodies with enhanced Fc-mediated effector functions [4, 5]. Although the results of this vaccine trial continue to be debated [6], particularly in light of the failure of the similarly designed HVTN 702 trial [7], these findings have led to the hypothesis that Fc-mediated antibody effector functions may afford some protection against HIV-1 in the absence of detectable neutralization.

Attempts to model the types of immune responses that may have contributed to protection in the RV144 trial have yielded mixed results. While a few prime-boost vaccine studies in macaques using different vectors, adjuvants and challenge viruses observed correlations of V2-specific antibodies with protection [8–13], others found no such association [14–18]. Passive administration of a V2-specific antibody with limited neutralization potency reduced viral loads in macaques following mucosal challenge with the tier 1 SHIV BaL.P4 [19]. Partial protection against mucosal SIV_mac_251 challenge was also achieved following the infusion of non-neutralizing IgG purified from vaccinated macaques with polyfunctional antibody signatures [20]. Furthermore, adeno-associated virus delivery of an antibody that directs ADCC against SIV-infected cells but lacks detectable neutralizing activity protected a single animal from mucosal SIV challenge [21]. However, the majority of passive transfer experiments with non-neutralizing antibodies (nnAbs) have failed to protect against SIV or SHIV challenge [22–24]. Thus, definitive evidence for protection by nnAbs has been elusive.

We and others have shown that ADCC correlates with neutralization and that nnAbs generally do not mediate ADCC against HIV- or SIV-infected cells [25–30]. This implies that most antibodies that are capable of binding to functional Env trimers on virions to block viral infectivity are also capable of binding to Env on the surface of virus-infected cells to mediate ADCC. Nevertheless, ADCC in the absence of detectable neutralization and neutralization in the absence of ADCC have been observed indicating that these antiviral functions may be uncoupled for certain antibodies [25, 30].

One such antibody showing an uncoupling of neutralization and ADCC for SIV_mac_239 is PGT145. This antibody binds to a proteoglycan epitope at the apex of HIV-1 Env trimers and potently neutralizes genetically diverse HIV-1 isolates [31]. We previously showed that PGT145 also binds to Env on the surface of SIV-infected cells and directs efficient NK cell killing of SIV-infected cells but only very weakly neutralizes SIV infectivity [32]. We proposed that the affinity of PGT145 for SIV Env is sufficient to allow enough binding to virus-infected cells to promote ADCC but not enough for potent neutralization. Indeed, a single amino acid substitution in the V2 core epitope of SIV Env increases PGT145 binding and confers notable sensitivity to neutralization [32]. In the present study, we took advantage of the functional properties of PGT145 in the context of SIV_mac_239 to investigate the extent to which ADCC is sufficient for protection and the effect of a minimal change in Env that makes the virus sensitive to neutralization.

## Results

Neutralization of HIV-1 by PGT145 is dependent on antibody binding to a quaternary proteoglycan epitope at the V2 apex that includes an N-linked glycan at position 160 (N160) and a polybasic region at the convergence of all three gp120 protomers [31]. PGT145 is therefore specific for Env trimers and does not bind to monomeric gp120 [33]. Using an assay designed to measure ADCC against virus-infected cells expressing natural conformations of Env [34], we found that PGT145 efficiently directs ADCC against SIV_mac_239-infected cells (**Fig. 1A**), but only weakly neutralizes replication-competent SIV_mac_239 at high concentrations (≥25 µg/ml) (**Fig. 1B**) [32]. The cross-reactivity of PGT145 with SIV is a consequence of the binding of this antibody to a conserved epitope at the V2 apex of SIV Env trimers that includes an N-linked glycan at position 171 (N171) and basic residues in the V2 core epitope [32]. Further analysis of Env substitutions in this region identified a single amino acid substitution (K180S) that enhances PGT145 binding to Env, increases the susceptibility of SIV-infected cells to ADCC (**Fig. 1A**), and is sufficient to confer sensitivity to neutralization (**Fig. 1B**) [32].

**Figure 1.**
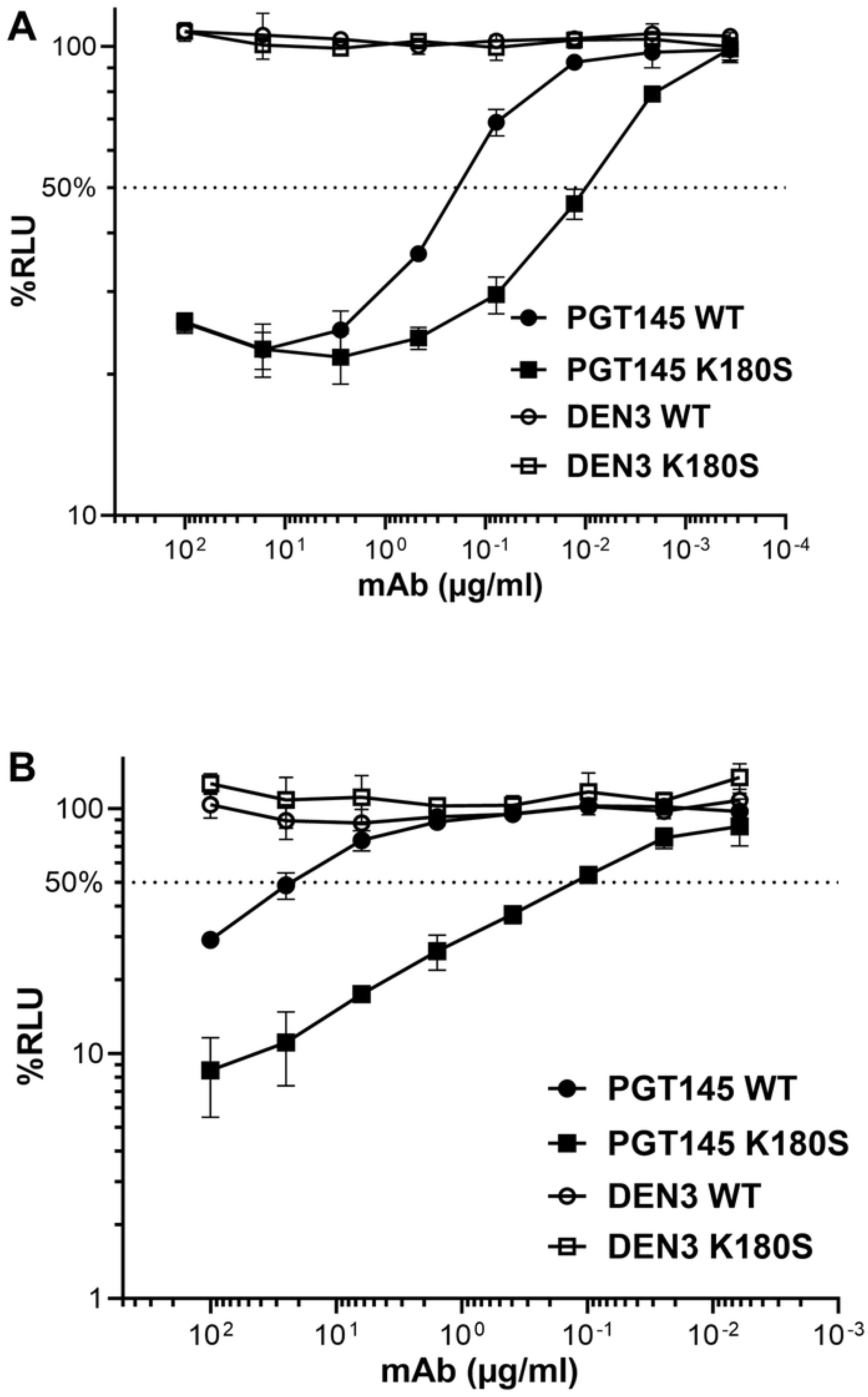
Sensitivity of SIV_mac_239 and SIV_mac_239 K180S to PGT145-mediated ADCC and neutralization. (A) CEM.NKR-_CCR5_-sLTR-Luc cells were infected with wild-type SIV_mac_239 (WT) or SIV_mac_239 K180S (K180S) and incubated with an NK cell line (KHYG-1 cells) expressing rhesus macaque CD16 at a 10:1 effector to target (E:T) ratio for eight hours in the presence of serial dilutions of mAbs PGT145 and DEN3. ADCC was calculated as the percent relative light units (%RLU) of luciferase activity remaining in SIV-infected cells incubated with antibody relative to SIV-infected cells without antibody after correcting for the background luciferase activity present in uninfected cells. (B) Wild-type SIV_mac_239 (WT) and SIV_mac_239 K180S (K180S) were incubated with serial dilutions of PGT145 and DEN3 for one hour before addition to TZM-bl cells. Neutralization was measured as the dose-dependent reduction in luciferase activity (% RLU) for viruses incubated with antibody relative to viruses without antibody after subtracting the background luciferase activity present in uninfected cells. ADCC and neutralization values reflect the mean and standard deviation (error bars) for triplicate wells at each antibody concentration. The dotted lines indicate half-maximal responses.

### PGT145 does not protect rhesus macaques against wild-type SIV_mac_239

To determine if ADCC is sufficient for protection, separate groups of six rhesus macaques of Indian ancestry were treated with PGT145 or a control antibody (DEN3). PGT145 and DEN3 were administered intravenously at doses of 30 mg/kg. After a five-day interval to allow the antibodies time to become distributed throughout the tissues [35], the animals were challenged intrarectally with an animal-titered stock of SIV_mac_239 at a dose sufficient to infect all the control animals (6,000 TCID_50_). On the day of challenge, the average concentration of PGT145 in plasma for the animals that received this antibody was 307 ± 58.5 µg/ml (**Table 1**). Plasma from each of the PGT145-treated animals collected on the day of challenge also mediated ADCC against SIV_mac_239-infected cells with a mean 50% ADCC titer of 239 ± 50 (**Fig. 2A** **and Table 1**). Consistent with only weak neutralization of wild-type SIV at high concentrations of PGT145, neutralizing antibodies to SIV_mac_239 were not detectable in serum from any of the animals (**Fig. 2B**). However, as expected these samples did neutralize SIVmac239 K180S (**Table 1**). Following intrarectal challenge, all of the animals became infected and viral loads did not differ significantly during acute or chronic infection between PGT145- and DEN3-treated animals (**Fig. 2C**). Thus, the potent ADCC of PGT145 against SIV-infected cells as measured in a standard in vitro assay does not predict protection against SIV_mac_239 and suggests that this antiviral function alone is not sufficient for protection against viral challenge.

**Figure 2.**
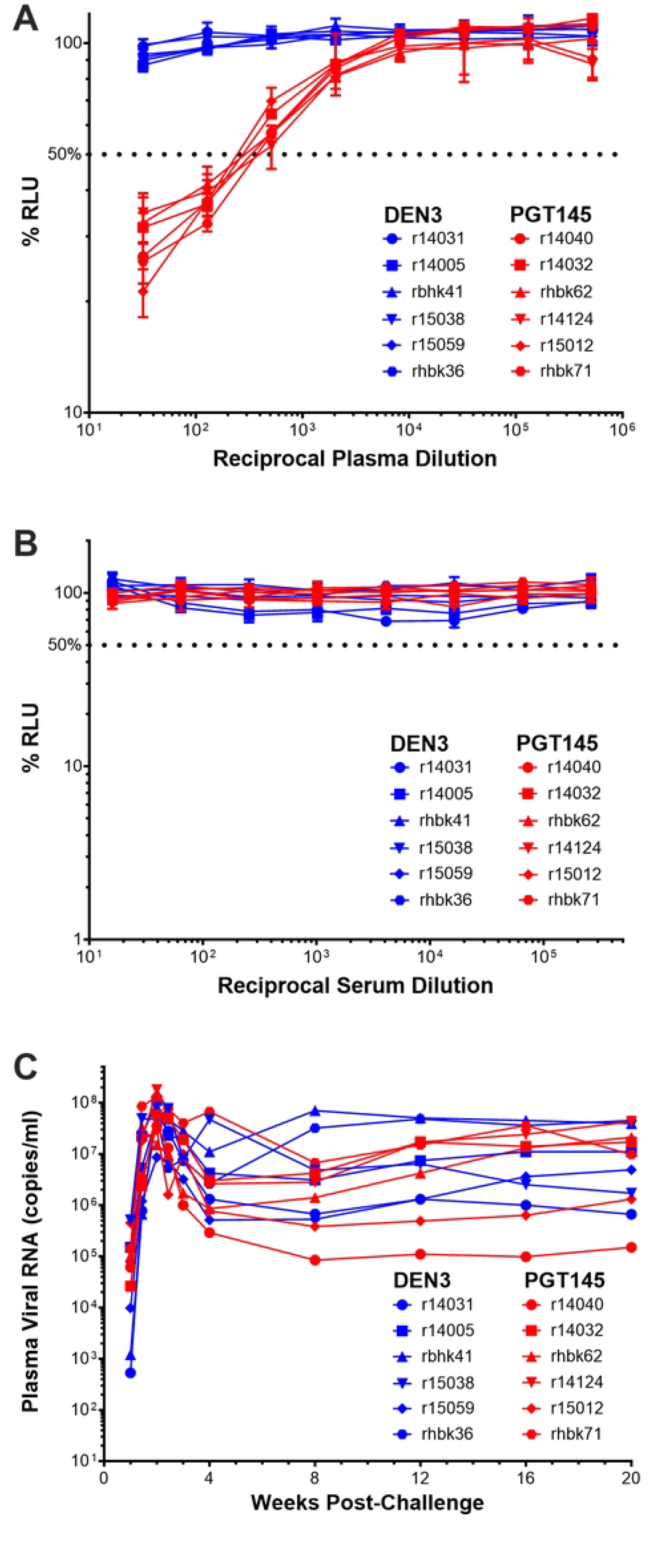
PGT145 does not protect against wild-type SIV_mac_239 challenge. (A) ADCC responses in plasma at the time of challenge. CEM._NKR-_CCR5-sLTR-Luc cells infected with SIV_mac_239 were incubated with a NK cell line expressing rhesus macaque CD16 at a 10:1 effector/target ratio in the presence of serial dilutions of plasma collected on the day of challenge from animals treated with either PGT145 or DEN3. (B) Neutralization activity in plasma at the time of challenge. ADCC was calculated from the dose-dependent loss of luciferase activity (% RLU) after an 8 hour incubation. SIV_mac_239 was incubated in the presence of serial dilutions of plasma for 1 hour before addition to TZM-bl cells. Luciferase induction was measured after 48 hours and neutralization was calculated from the dose-dependent inhibition of luciferase (% RLU). (C) Post-challenge viral loads. Viral loads in plasma were measured using a qRT-PCR assay with a detection threshold of 15 copies/ml.

**Table 1.** PGT145, ADCC and neutralizing antibody titers on the day of challenge.

| Study | Animal | PGT145 (µg/ml) | 50% ADCC titer<sup>a</sup> | 50% Neut. titer<sup>b</sup> |
| --- | --- | --- | --- | --- |
| Wt 239 challenge | r14040 | 367 | 245 | 695 |
|  | r14032 | 340 | 201 | 483 |
|  | rhbk62 | 319 | 217 | 461 |
|  | r14124 | 293 | 313 | 850 |
|  | r15012 | 214 | 181 | 927 |
|  | rhbk71 | - | 279 | 466 |
|  | Mean ± SD | 307 ± 58.5 | 239 ± 50.0 | 647 ± 208 |
| K180S challenge | rhbj64 | 330 | 434 | 454 |
|  | rh2761 | 280 | 398 | 725 |
|  | rhbj89 | 280 | 888 | 1251 |
|  | r09070 | 395 | 452 | 685 |
|  | r14054 | 252 | 1406 | 1205 |
|  | r15003 | 243 | 1399 | 1571 |
|  | Mean ± SD | 296 ± 57.1 | 830 ± 479 | 982 ± 425 |

<sup>a</sup>ADCC responses were measured against SIV<sub>mac</sub>239-infected cells for the animals challenged with wild-type SIV<sub>mac</sub>239 and against SIV<sub>mac</sub>239 K180S-infected cells for the animals challenged with SIV<sub>mac</sub>239 K180S.
<sup>b</sup>Neutralizing antibody titers were measured against SIV<sub>mac</sub>239 K180S for the animals challenged with both wild-type SIV<sub>mac</sub>239 and SIV<sub>mac</sub>239 K180S.

### PGT145 partially protects rhesus macaques against SIV_mac_239 K180S

We previously demonstrated that the relatively low binding affinity of PGT145 for SIV Env is sufficient to trigger ADCC by cross-linking multiple Env trimers on the surface of SIV_mac_239-infected cells to Fcγ receptors on NK cells, but is not high enough to block viral infectivity [32]. However, a single amino acid substitution in the V2 core epitope (K180S) increases PGT145 binding to Env and ADCC by approximately 100-fold [32]. This substitution also renders SIV_mac_239 K180S susceptible to neutralization by PGT145 (**Fig. 1B**). To determine if the combined antiviral effects of neutralization and ADCC translate into better protection, PGT145 and DEN3 were administered intravenously to two additional groups of six rhesus macaques at doses of 30 mg/kg and protection was assessed five days later by intrarectal challenge with SIV_mac_239 K180S. The challenge dose for this study was increased five-fold to 30,000 TCID_50_, as this was determined to be the minimum dose required to infect three of three naïve macaques by intrarectal inoculation in a prior animal titration experiment.

On the day of challenge, the average PGT145 concentration in plasma among the animals that received this antibody was 296 ± 57.1 µg/ml (**Table 1**). Potent ADCC and neutralizing antibody responses to SIV_mac_239 K180S were detectable with mean 50% titers of 830 ± 479 and 982 ± 425, respectively (**Fig. 3A, 3B and Table 1**). PGT145 was also detectable in rectal transudate at 0.90-6.3% and 1.3-4.0% (median 4.2% and 2.8%) of total IgG confirming that the antibody was present at the site of challenge (**Fig. 3C**). Following SIV_mac_239 K180S challenge, five of the six animals in each group became infected (**Fig. 3D**). PGT145 therefore did not prevent virus acquisition, even at a challenge dose that was not quite high enough to infect all the control animals. However, significant differences in the timing and peak of viremia were observed. Compared to the DEN3-treated control group, peak viral RNA loads in plasma during acute infection were delayed by 1.4 weeks (p=0.0004, SE=0.24, linear regression) and were 1.28 log lower (p=0.0006, SE=0.24, linear regression) in the PGT145-treated animals (**Fig 3C**). During chronic infection (8-24 weeks post-challenge) viral loads were also significantly lower in the PGT145-treated animals (p=0.008, SE=0.7, linear mixed models) (**Fig. 3C**). Therefore, even though PGT145 did not prevent SIV_mac_239 K180S transmission, these viral load differences reveal a significant impact of the antibody on virus replication.

**Figure 3.**
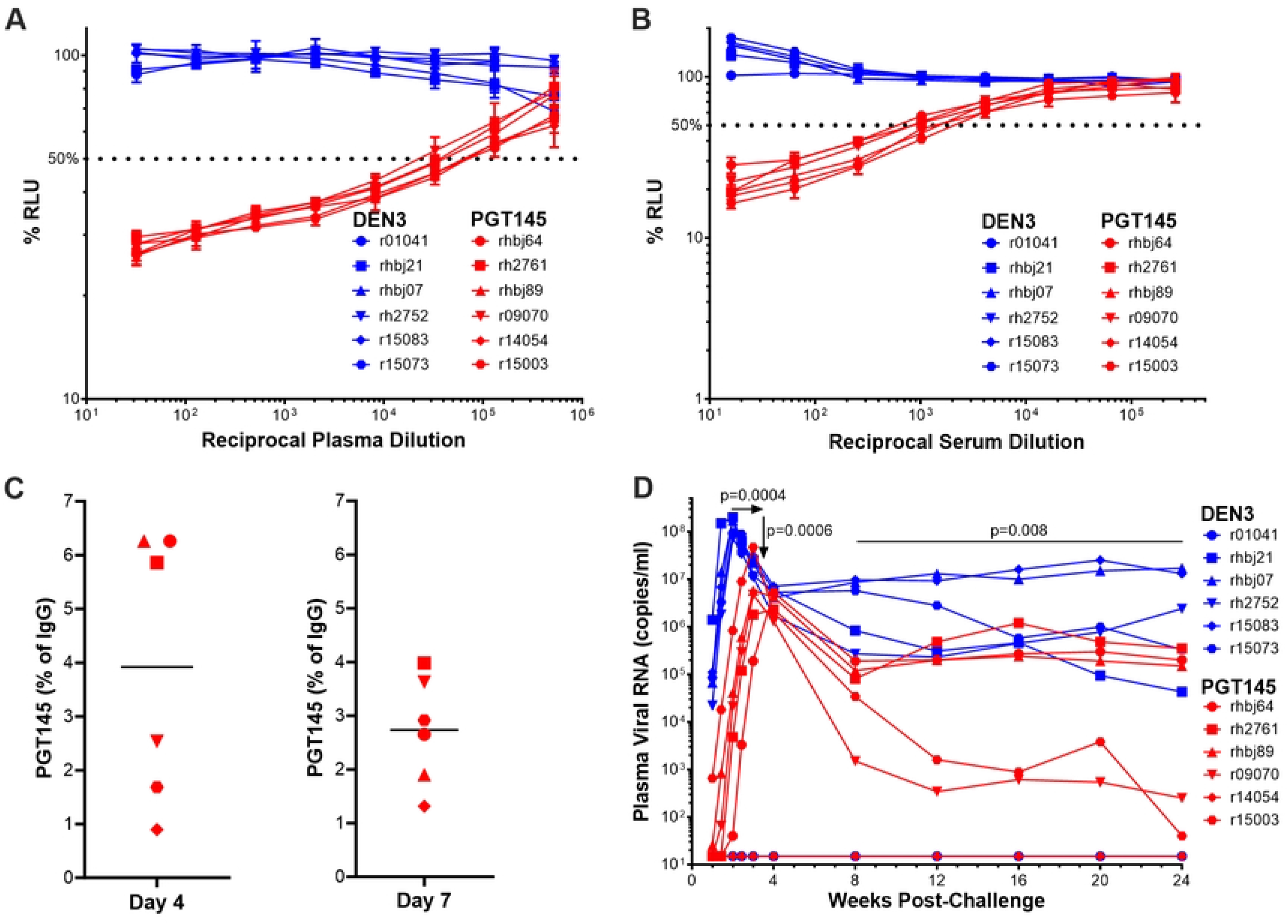
PGT145 affords partial protection against SIV_mac_239 K180S challenge. (A) ADCC responses in plasma at the time of challenge. CEM._NKR-_CCR5-sLTR-Luc cells infected with SIV_mac_239 K180S were incubated with a NK cell line expressing rhesus macaque CD16 at a 10:1 effector/target ratio in the presence of serial dilutions of plasma collected on the day of challenge from animals treated with either PGT145 or DEN3. ADCC was calculated from the dose-dependent loss of luciferase activity (% RLU) after an 8 hour incubation. (B) Neutralization activity in serum at the time of challenge. SIV_mac_239 K180S was incubated in the presence of serial dilutions of serum for 1 hour before addition to TZM-bl cells. Luciferase induction was measured after 48 hours and neutralization was calculated from the dose-dependent inhibition of luciferase (% RLU). (C) PGT145 concentrations as a percentage of total IgG in rectal transudate. PGT145 and total IgG concentrations were measured by ELISA in mucosal secretions eluted from rectal swabs collected on days 4 and 7 post-challenge. (D) Post-challenge viral loads. Viral loads in plasma were measured using a qRT-PCR assay with a detection threshold of 15 copies/ml. Statistics were calculated using a mixed effects model by comparing results from acute (weeks 1-4) and chronic (weeks 8-24) infection to pre-infection (week 0).

Nevertheless, the protection afforded by PGT145 was not as complete as might have been expected based on a previous meta-analysis of the relationship between neutralizing antibody titers and protection in previous passive immunization studies [36]. The mean 50% neutralization titer (ID50) in serum against SIV_mac_239 K180S on the day of challenge was 982 ± 425, which exceeded the ID50 titer of 685 (95% CI 319, 1471) predicted to afford 95% protection against mucosal SHIV challenge [36]. The explanation for this difference is presently unclear but may reflect resistance of the challenge virus to complete neutralization by PGT145. To investigate this possibility, serial dilutions of the SIV_mac_239 K180S challenge stock were incubated with 50 µg/ml of PGT145 and K11 before the addition of TZM-bl cells. In contrast to K11, which binds to a glycan hole on the gp120 surface of the SIV Env trimer and completely neutralizes SIV_mac_239 K180S at high multiplicities of infection [37], residual infectivity became detectable at a 10-fold virus dilution (1.4 ng/ml p27) in the presence of PGT145 (**S1 Fig.**). These results are similar to the previously reported incomplete neutralization of HIV-1 by V2 apex bnAbs [38, 39]. The limited protection against SIV_mac_239 K180S challenge may therefore be due to a fraction of the challenge virus that is resistant to PGT145, possibly as a consequence of heterogenous Env glycosylation.

### PGT145 selects for escape mutations in Env

To better understand incomplete protection against SIV_mac_239 K180S, the virus population in plasma was sequenced at weeks 4 and 8 post-challenge. These time points were selected to capture the emergence of antibody escape mutations when virus replication was occurring in the presence of passively administered PGT145 (**S2 Fig.**). While the K180S change was retained as a fixed substitution in all the animals at both time points, several Env changes were observed in two or more of the PGT145-treated animals that were not observed in the DEN3-treated control animals. With a couple of exceptions, these Env substitutions were present at a low frequency that did not exceed 75% of the virus population (**Fig. 4A**). The low frequency of these substitutions probably reflects the incomplete and transient nature of PGT145 selection, owing to a fraction of neutralization-resistant virus in the challenge stock (**S1 Fig.**) coupled with declining concentrations of PGT145 in serum after passive transfer (**S2 Fig.**).

**Figure 4.**
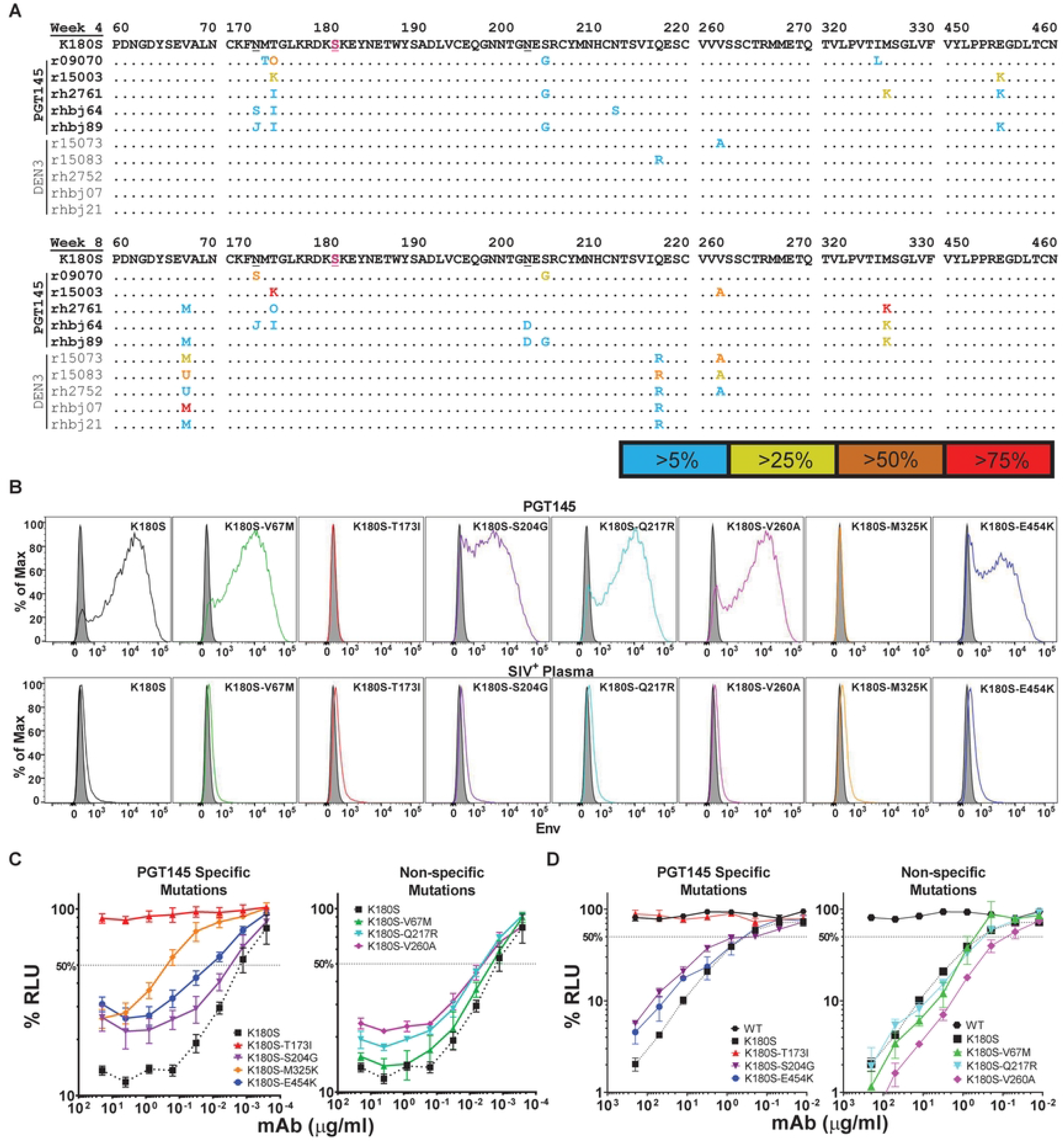
PGT145 treatment selects for amino acid changes in Env in vivo that confer resistance to ADCC and to neutralization. (A) Viral RNA was isolated from plasma at weeks 4 and 8 post-infection and sequenced. The predicted amino acid sequences in Env from PGT145-treated animals (top, bold) and from DEN3-treated animals (bottom, plain text) were aligned to SIV_mac_239 K180S Env. Regions of Env with substitutions in multiple animals are shown. The K180S substitution is indicated in magenta and residues N171 and N202 are underlined in the SIV_mac_239 K180S Env reference sequence. Positions of identity are indicated by periods and amino acid differences are identified by their single letter code. Amino acid ambiguities are indicated with non-standard letters as follows: U = M/L, J = S/K, O = A/I (week 4) and O = I/K (week 8). The frequencies of each substitution within the virus population are indicated by color. (B) Antibody binding to Env was assessed by staining CEM._NKR-_CCR5-sLTR-Luc cells infected with each of the indicated SIV_mac_239 K180S mutants with PGT145 (top row) and with SIV+ plasma (bottom row) followed by AF647-conjugated goat anti-human IgG F(ab′)_2_ after gating on virus-infected (CD4_low_, p27^+^) cells. The shaded histograms represent non-specific staining with DEN3. (B & C) SIV_mac_239 K180S mutants with substitutions in Env selected only in PGT145-treated animals (specific) or in both PGT145- and DEN3-treated animals (non-specific) were tested for susceptibility to ADCC (B) and neutralization (C) mediated by PGT145. The dotted lines indicate half-maximal ADCC or neutralization and the error bars indicate standard deviation of the mean for triplicate wells at each antibody concentration.

At week 4 post-infection, the virus population of all five of the PGT145-treated animals acquired changes in residues 171-173 that coincide with the loss of an N-linked glycosylation site previously shown to be essential for PGT145 binding (**Fig. 4A**) [32]. Two additional substitutions were also detected in three of these animals (S204G and E454K). The S204G substitution disrupts another N-linked glycosylation site (residues 202-204) that was not previously implicated in Env binding by PGT145. At week 8 post-infection, substitutions in residues 171-173 persisted in four of the PGT145-treated animals and increased in frequency in two of them (**Fig. 4A**). Amino acid changes in residues 202-204 were also present in three animals (**Fig. 4A**). Although the E454K substitution was not detected at week 8, the virus acquired another substitution (M325K) in three of the PGT145-treated animals (**Fig. 4A**). Additional Env changes unrelated to selection by PGT145 were also observed in the DEN3-treated animals (V67M, Q217R and V260A) that may be related to the adaptation of SIV_mac_239 K180S for more efficient replication in macaques (**Fig. 4A**).

Env substitutions selected in SIV-infected macaques were introduced into SIV_mac_239 K180S to assess their impact on the sensitivity of virus-infected cells to PGT145 binding and ADCC. These included four changes that emerged in the PGT145-treated animals (T173I, S204G, M325K and E454K) and three in the DEN3-treated control animals (V67M, Q217R and V260A). None of these substitutions impaired Env expression on the surface of infected cells as indicated by similar levels of Env staining using plasma pooled from SIV-infected animals (**Fig. 4B**). The V67M, Q217R and V260A substitutions that arose in the control animals also did not reduce PGT145 staining (**Fig. 4B**). In contrast, each of the substitutions selected in the PGT145-treated animals partially or completely impaired PGT145 binding to virus-infected cells. PGT145 staining was diminished by S204G and E454K and nearly eliminated by T173I and M325K (**Fig. 4B**). These differences in Env binding corresponded to differences in the susceptibility of virus-infected cells to ADCC. Whereas the Env changes acquired in the control animals had little effect on sensitivity to PGT145, the substitutions selected in the PGT145-treated animals conferred partial or complete resistance to ADCC (**Fig. 4C**). In accordance with their effects on Env binding, sensitivity to ADCC was impaired to a greater extent by T173I and M325K than by S204G or E454K (**Fig. 4C**).

Similar effects were observed on the sensitivity of each of the Env variants to neutralization. The Env substitutions that arose in the DEN3-treated control animals did not alter (V67M & Q217R) or slightly increased (V260A) the sensitivity of SIV_mac_239 K180S to neutralization by PGT145 (**Fig. 4D**). In contrast, the substitutions selected in the PGT145-treated animals conferred partial (S204G and E454K) or complete (T173I) resistance to neutralization (**Fig. 4D**). The T173I substitution is therefore sufficient for complete resistance to both neutralization and ADCC consistent with the requirement for glycosylation of N171 for PGT145 binding [32].

### Structural analysis of PGT145 escape mutations

Two cryo-electron microscopy structures of the SIV_mac_239 Env trimer were recently solved [37, 40], one of which was determined for PGT145 in complex with an Env variant (SIV_mac_239 Env:K180T) with a lysine-to-threonine (K180T) substitution that stabilizes the binding of this antibody to SIV Env similar to our K180S substitution [32, 40]. This structure reveals extensive asymmetric contacts between PGT145 and the N171 glycans on each of the three gp120 protomers, primarily involving the complementarity determining region 3 and 1 of the heavy and light chain, respectively (CDRH3 and CDRL1) (**Fig. 5A**). This is consistent with the complete resistance of the T173I variant of SIV_mac_239 K180S to ADCC and neutralization as a result of the loss of these N-linked glycans (**Fig. 4**). PGT145 is also near another glycan attached to residue N202 (**Fig. 5A**). The N202 glycans are located on the periphery of the antibody binding site at a distance of 6-10 Å from the PGT145 light chain and may have a greater role in the coordination of antibody binding rather than forming direct protein contacts. This could account for the modest effect of the loss of these glycans on the sensitivity of the S204G variant to ADCC and neutralization (**Fig. 4**).

**Figure 5.**
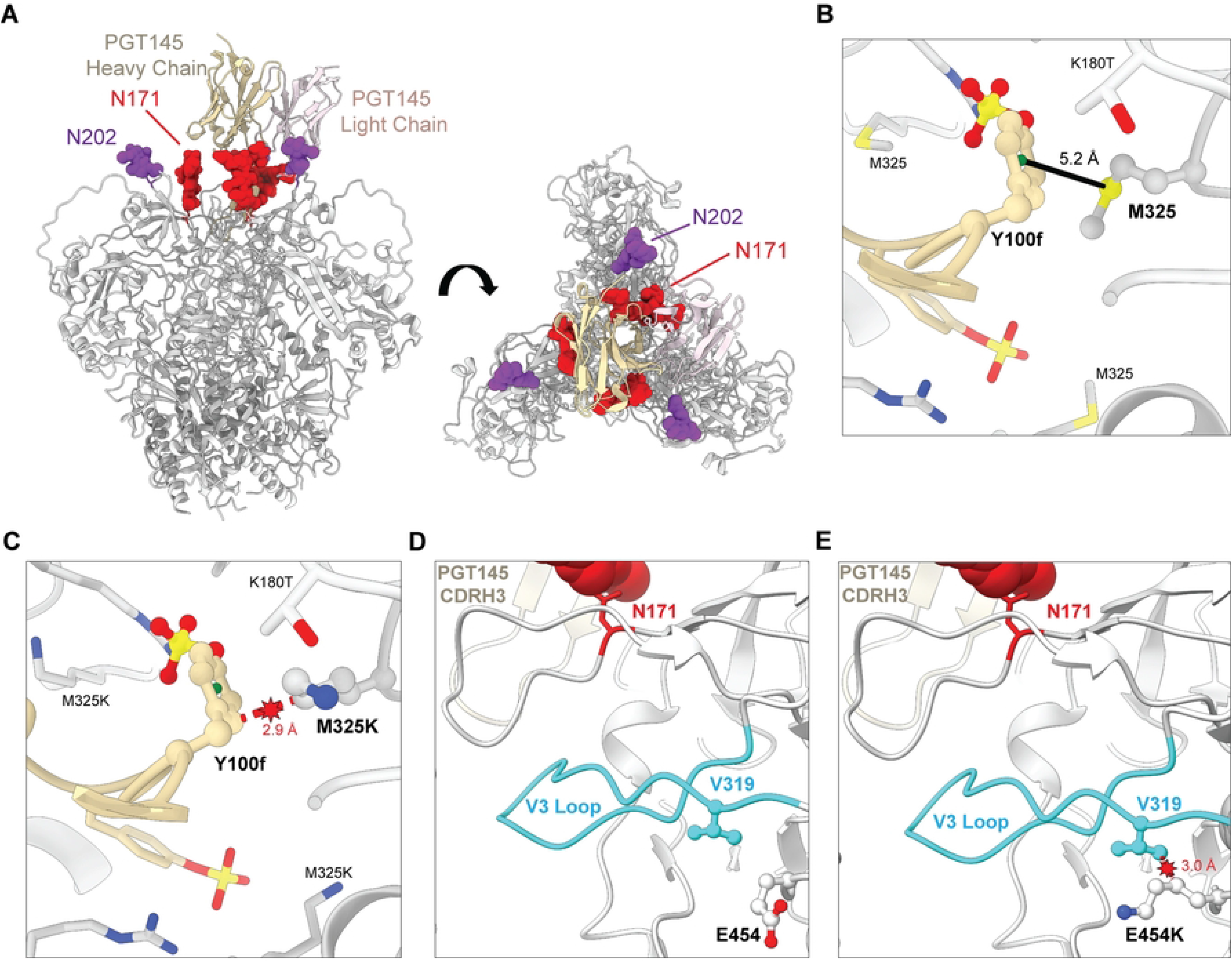
Structural analysis of Env substitutions selected in SIV_mac_239 K180S. The amino acid changes that emerged in PGT145-treated animals infected with SIV_mac_239 K180S were mapped onto a cryo-EM structure of PGT145 in complex with the SIV_mac_239:K180T Env trimer (PDB:8DVD) [40]. (A) Orthogonal views parallel (left) and perpendicular (right) to the viral membrane are shown for the variable heavy (tan) and light (pink) chains of PGT145 bound to the V2 apex of the SIV_mac_239:K180T Env trimer (gray). The N-linked glycans on residues N171 (red) and N202 (purple) are indicated. (B & C) An interaction between M325 of SIV Env and the aromatic ring of Y100f in the PGT145 VH chain contributes to PGT145-Env binding (B) and M325K is predicted to clash with Y100f of the PGT145 VH chain and K180T of SIV Env (C). (D & E) E454 is not part of the PGT145 binding interface (D). However, the E454K substitution may interfere allosterically with PGT145 binding by clashing with V319 and altering the V3 loop (light blue) (E). Three-dimensional models were generated using ChimeraX (ver 1.5) [51].

The methionine at position 325 of SIV Env also participates directly in PGT145 binding. M325 is located in the V3 loop within the core of the Env trimer where it contacts a sulfated tyrosine residue (Y100f) of the PGT145 CDRH3 (**Fig. 5B**) [40]. M325 and Y100f form a characteristic methionine-aromatic interaction that is predicted to contribute an additional 1-1.5 kcal/mol of energy to protein stability beyond a purely hydrophobic interaction [41]. Modeling of the M325K substitution in the SIV Env trimer suggests that a lysine at this position would disrupt the interaction with PGT145 by clashing with Y100f (**Fig. 5C**), which would account for the decreased sensitivity of the M325K variant to PGT145 binding and ADCC (**Fig. 4B and 4C**).

The glutamic acid at position 454 of SIV Env is distal to the PGT145 interface, so the E454K substitution is unlikely to alter direct contacts with PGT145 (**Fig. 5D**). However, a non-conservative change to a lysine residue at this position with a bulkier side chain is predicted to clash with V319 on the V3 loop, which may have allosteric effects on PGT145 binding (**Fig. 5E**). Accordingly, such allosteric effects may account for the diminished binding and ADCC activity of PGT145 against cells infected with the E454K variant of SIV_mac_239 K180S (**Fig. 4B and 4C**).

## Discussion

Although neutralizing antibodies have repeatedly demonstrated protection against SHIVs in nonhuman primates [36], definitive evidence of protection by non-neutralizing antibodies has been elusive. To determine the extent to which neutralizing and non-neutralizing activities are required for protection against SIV, we took advantage of the unique cross-reactivity of the HIV-1 bnAb PGT145 with the SIV envelope glycoprotein. PGT145 binds to Env on the surface of SIV-infected cells and directs efficient NK cell killing of virus-infected cells, but poorly neutralizes SIV infectivity. Despite mediating potent ADCC both *in vitro* and *ex vivo*, this antibody failed to prevent mucosal transmission or reduce virus replication in macaques after intrarectal challenge with wild-type SIV_mac_239. However, PGT145 was able to significantly delay and reduce virus replication in animals challenged with a variant of SIV_mac_239 that contains a single amino acid substitution that increases its binding to Env and renders the virus susceptible to neutralization [32].

The inability of PGT145 to protect against wild-type SIV_mac_239 challenge despite potent ADCC against this virus raises questions about the extent to which ADCC alone is sufficient to prevent immunodeficiency virus transmission. The ADCC assay used here is the same assay that we used to measure ADCC against HIV-infected cells in the immune correlates analysis of the RV144 trial [2]. The ADCC titers in our animals were much higher than ADCC titers in RV144 vaccine recipients, which were often undetectable and never exceeded 35% killing at the highest serum concentration tested [2]. Thus, the absence of protection against wild-type SIV_mac_239 cannot simply be explained by suboptimal ADCC responses. There are of course important differences that limit the extent to which the results of this study may be compared with clinical vaccine trials. The passive administration of a single monoclonal antibody does not represent the breadth of polyclonal antibodies elicited in response to vaccination. Our animals were also challenged with a sufficiently high dose of SIV_mac_239 to establish infection in enough control animals that the outcome of protection would be interpretable after a single intrarectal inoculation. It is conceivable that this challenge dose may have overwhelmed partial protection that might have been observed at a more physiological challenge dose. However, the absence of any detectable effect on post-challenge viral loads after SIV_mac_239 transmission suggests otherwise.

PGT145 did however confer partial protection against SIV_mac_239 K180S, as indicated by a delay in peak viremia as well as significant reductions in viral loads during acute and chronic infection. Since the K180S substitution increases PGT145 binding to Env by approximately 100-fold and confers sensitivity to neutralization [32], these results are consistent with studies showing that neutralization is a key determinant of antibody-mediated protection [36, 42, 43]. However, the protection afforded by PGT145 was not as complete as might have been expected based on a meta-analysis of the relationship between neutralizing antibody titers and protection in previous passive immunization studies [36]. On the day of challenge, the mean ID50 titer in serum against SIV_mac_239 K180S (982 ± 425) exceeded the ID50 titer of bnAbs predicted to achieve 95% protection against mucosal SHIV challenge (685, 95% CI 319, 1471) [36]. The explanation for this difference is not certain but may reflect a fraction of the challenge virus that is resistant to complete neutralization by PGT145 as a consequence of heterogenous Env glycosylation (**S2 Fig.**). Indeed, this possibility is supported by previous reports of incomplete neutralization of HIV-1 by V2 apex bnAbs [38, 39].

The rapid emergence of amino acid changes in Env that further confer resistance to PGT145 helps to explain breakthrough replication of SIV_mac_239 K180S in the presence of high *in vivo* antibody concentrations. Substitutions that disrupt a conserved N-linked glycosylation site at position 171 were observed in each of the PGT145-treated animals. We previously demonstrated that glycosylation of residue N171 is essential for PGT145 binding to the SIV Env trimer [32] and that this site corresponds to an N-linked glycan at position 160 of HIV-1 Env (N160) that is part of the V2 epitope for this antibody [31]. Extensive contacts between PGT145 and the glycans attached to N171 on each of the gp120 protomers were confirmed in a recent cryo-EM structure of PGT145 bound to the SIV_mac_239 K180T Env trimer [40]. Accordingly, a threonine-to-isoleucine substitution at residue 173 (T173I), which arose in four of the five animals by week 4 post-infection, completely abrogated the sensitivity of SIV_mac_239 K180S to neutralization and ADCC. A serine-to-glycine substitution at position 204 (S204G) that is predicted to eliminate another N-linked glycan attached to residue N202 also impaired the susceptibility of SIV_mac_239 K180S to ADCC and neutralization. This glycan is located at the periphery of the PGT145 binding site and does not appear to make direct contact with the antibody, which may account for the more modest effect of the loss of this glycan on PGT145 binding, ADCC, and neutralization compared to T173I [40].

Additional Env changes selected in the PGT145-treated animals may also have contributed to viral breakthrough. A methionine-to-lysine substitution at position 325 (M325K) emerged in three animals by eight weeks post-challenge that disrupts a methionine-aromatic interaction between M325 and a sulfated tyrosine residue (Y100f) in the HCDR3 region of PGT145 [40]. Consistent with the replacement of this stabilizing interaction with a predicted clash between these two residues, the M325K substitution reduced the sensitivity of virus-infected cells to PGT145 binding and ADCC by more than two orders of magnitude. The glutamic acid-to-lysine substitution at position 454 (E454K), which was present in three of the animals at week four but had disappeared by week eight post-challenge, is unlikely to affect PGT145 binding directly since it is located distal to the antibody binding site [40]. However, this substitution may have allosteric effects on Env as a result of conformational changes to the V3 loop that could account for diminished PGT145 binding and ADCC responses to virus-infected cells.

Overall, this study shows that in the case of a V2-specific antibody with potent ADCC against immunodeficiency virus-infected cells, that ADCC in the absence of detectable neutralization is not sufficient for protection. However, partial protection may be achieved by increasing the affinity of antibody binding to Env above the threshold required for neutralization of viral infectivity. Therefore, while ADCC and potentially other Fc-mediated effector functions may contribute to protection through the elimination of productively infected cells, the higher affinity of Env binding necessary for potent neutralization is a critical determinant of antibody-mediated protection.

## Materials and Methods

### Ethics Statement

Twenty-four rhesus macaques (*Macaca mulatta*) of Indian ancestry were used in this project. These animals were housed at the Wisconsin National Primate Research Center (WNPRC) in accordance with the standards of the Association for the Assessment and Accreditation of Laboratory Animal Care (AAALAC) and the University of Wisconsin Research Animal Resources Center and Compliance unit (UWRARC). Animal experiments were approved by the University of Wisconsin College of Letters and Sciences and the Vice Chancellor for Research and Graduate Education Centers AICUC (protocol number G005141) and performed in compliance with the principles described in the *Guide for the Care and Use of Laboratory Animals* [44]. Fresh water was always available, commercial monkey chow was provided twice a day and fresh produce was supplied daily. To minimize any pain and distress related to experimental procedures, Ketamine HCL was used to sedate animals prior to blood collection and animals were monitored twice a day by animal care and veterinary staff. Animals positive for MHC class I alleles associated with spontaneous control of SIV infection (*Mamu-B*008* and *-B*017*) were excluded and experimental and control groups were structured with an equal distribution of male and female animals.

### Antibody infusion

Purified, low-endotoxin stocks of PGT145 and DEN3 were produced by Catalent Pharma Solutions (Madison, WI) and The Scripps Research Institute (La Jolla, CA), respectively. These antibodies were diluted in sterile phosphate buffered saline (PBS) and gradually administered to rhesus macaques at doses of 30 mg/kg by intravenous infusion through a catheter placed aseptically in the saphenous vein of anesthetized animals over a period of approximately 20-30 minutes to prevent anaphylaxis.

### Antibody concentrations in plasma and rectal transudate

Rectal mucosal secretions were collected and processed with a modified wick method using Weck-Cel spears (Beaver Visitec, # 0008680) as described previously [45]. Total IgG content was determined by ELISA using purified mAbs PGT121 or PGT145 as a standard. Plates were coated with 2 µg/ml anti-human F(ab’)2-specific antibody (Jackson ImmunoResearch Laboratories #109-005-097) and blocked using PBS containing 3% BSA. Sera and rectal samples were diluted 1:200 or 1:50 in 1% BSA, respectively, and serially diluted in PBS/1% BSA. Antibodies were allowed to bind for 1 hour at room temperature before plates were washed 3x with PBS/0.05% Tween 20. Bound antibodies were then detected using 1:2000 diluted alkaline phosphatase-labelled anti-human IgG Fcγ (Jackson ImmunoResearch Laboratories #109-055-098) and 4-Nitrophenyl phosphate disodium salt hexahydrate substrate (Sigma #S0942). Optical density was measured at 405 nm and IgG concentrations were calculated based on the PGT121 or PGT145 standards.

For the detection of specific IgG, plates were coated with 2 µg/ml anti-His (Invitrogen #MA1-21315) over night and blocked for 1hour as described above. Soluble HIV-1 BG505 SOSIP trimers containing a hexahistidine tag were then allowed to bind at 3 µg/ml for 1hour before plates were washed 3 times with PBS/0.05%Tween 20. Serially diluted samples were then incubated for 1 hour. After washing 3 times with PBS/0.05%Tween 20, bound antibodies were detected as described above and specific antibodies concentrations calculated based on the PGT145 standard data.

### SIV challenge stocks and mucosal challenges

SIV challenge stocks were prepared by transfecting 293T cells with full-length infectious molecular clones of SIV_mac_239 and SIV_mac_239 K180S and then expanding the virus on activated rhesus macaque PBMCs. Cell culture supernatant was collected from infected rhesus macaque lymphocytes and virus yields were determined by SIV p27 ELISA and by qRT-PCR to determine SIV RNA copies/ml. Infectivity titers (TCID_50_/ml) were determined by limiting dilution on CEMx174 cells.

Rhesus macaques were inoculated intrarectally with SIV_mac_239 or SIV_mac_239 K180S. Thirty minutes prior to challenge, cryopreserved vials of each virus were thawed. Wild-type SIV_mac_239 was diluted to 6,000 TCID_50_ per ml and SIV_mac_239 K180S was diluted to 30,000 TCID_50_ per ml in serum-free RPMI and loaded into one ml tuberculin syringes. Animals were anesthetized, and with their pelvises elevated, the syringe (with the needle removed) was gently inserted into the rectum about 4 cm, then slightly withdrawn. The virus inoculum was slowly delivered to the rectal mucosa over about 1 minute. After withdrawing the syringe, it was examined for blood to ensure that there was no trauma to the rectum. To prevent drainage from the site of inoculation, the pelvic region of the animals was kept elevated for up to 30 minutes before returning them to their cages.

### Neutralization assay

Virus stocks for neutralization assays were prepared by transfecting 293T cells with full-length infectious molecular clones for SIV_mac_239 and SIV_mac_239 K180S and stored at -80°C. Neutralization of viral infectivity was measured using a standard TZM-bl reporter assay as previously described [46, 47]. TZM-bl cells were seeded at 5,000 cells per well (100 µl) in 96-well plates the day before the assay. SIVmac239 (2 ng p27) or SIVmac239 K180S (5 ng p27) were incubated with serial dilutions of antibody or serum (100 µl) for 1 h at 37°C before being added to TZM-bl reporter cells. After 3 days, luciferase activity in cell lysates was measured and virus neutralization was calculated from the reduction in RLU relative to cells incubated with virus but without antibody or serum. Additional wells plated with uninfected TZM-bl cells were also included to control for background luciferase activity.

### ADCC assay

To maximize infection of the target cells for measuring ADCC, VSV G-pseudotyped stocks of SIVmac239, SIVmac239 K180S and SHIVAD8-EO were prepared by co-transfecting 293T cells with *vif*-deleted clones for each virus together with a VSV G-expression construct and collecting supernatant 48 hours later. ADCC was measured as previously described [34, 48]. CEM.NKR-CCR5-sLTR-Luc cells, which express firefly luciferase under the control of the SIV LTR promoter, were inoculated with VSV G-pseudotyped SIVmac239, SIVmac239 K180S and SHIVAD8-EO. Two days later, the virus-infected target cells were incubated in triplicate with serial dilutions of antibody or plasma and an NK cell line (KHYG-1 cells) expressing rhesus macaque CD16 at a 10:1 effector-to-target ratio. Infected target cells incubated with NK cells but without antibody or plasma were included to determine maximal luciferase activity and uninfected target cells incubated with NK cells were included to determine background luciferase activity. ADCC was calculated as the percent relative light units (RLU) of luciferase remaining after an 8 hour incubation [% RLU = (experimental – background)/(maximal – background) X 100].

### Plasma viral loads

Plasma samples were isolated from blood collected with EDTA as an anticoagulant, cryopreserved at -80°C and SIV RNA levels were determined using a real-time RT PCR assay based on amplification of a conserved sequence in the SIV *gag* gene [49].

### Generation of SIV_mac_239 K180S mutants

Nucleotide substitutions were introduced into the p239SpE3’:K180S hemiviral plasmid by site-directed mutagenesis (New England Biolabs). Mutated p239SpE3’:K180S hemiviral plasmids were digested with SphI-XhoI and joined with the 5’ half of SIV_mac_239 SpX and SIV_mac_239 SpX ΔVif. Reconstructed full-length infectious molecular clones were sequence confirmed by the UW-Madison Biotechnology Center. Proviral DNA without the ΔVif mutation was transfected into HEK293T cells using GenJet (SignaGen) to generate viruses for neutralization assays. SIV_mac_239 SpX ΔVif plasmids were co-transfected with VSV-G envelope (pHDM.NJ strain) in a 2:1 ratio to generate viruses for ADCC assays and flow cytometry analyses. Culture supernatants were collected 48 h post-transfection, cleared of cell debris by centrifugation and concentrated on 50K MWCO centrifugal filters (Millipore Sigma). Concentrated viruses were well-mixed, aliquoted, and stored at -80°C. Concentrations were determined by anti-p27 ELISA (ABL, Inc.).

### Env staining

CEM.NKR-CCR5-sLTR-Luc cells were infected with *vif*-deleted SIVmac239, with or without Env mutations and pseudotyped with VSV G by spinoculation at 1200 g for 1 hour in the presence of 40 μg/ml Polybrene. Antibody binding to Env was evaluated by staining the cells 3 days post-infection. The cells were treated with Live/Dead NEAR IR viability dye (Invitrogen), washed in staining buffer (PBS with 1% FBS), and then incubated on ice for 30 minutes with 10 μg/ml of Env-specific antibody or a 1:32 dilution of SIV+ plasma pool from chronically infected rhesus macaques. A dengue virus-specific antibody (DEN3) was used as a negative control. An AF647-conjugated goat anti-human F(ab′)_2_ (Jackson Immunoresearch) and PE-Cy7-conjugated anti-CD4 (Clone OKT4, Biolegend) were used for subsequent staining on ice. For intracellular p27 staining, cells were fixed in PBS with 2% paraformaldehyde, washed in staining buffer, and stained with FITC-conjugated anti-SIV Gag antibody (clone 552F12) in Perm/Fix Medium B (Invitrogen). After washing, the cells were fixed in PBS with 2% paraformaldehyde and analyzed using a BD FACS Symphony flow cytometer. Antibody binding to Env was assessed by Env staining on the surface of SIV-infected (CD4_low_Gag^+^) cells.

### SIV sequencing

Replicating SIV was sequenced, as previously described [50]. Briefly, Qiagen MinElute virus spin kits were used to isolate vRNA. We then generated cDNA and amplified it using the SuperScript III one-step reverse-transcription (RT)-PCR with Platinum Taq High Fidelity, along with four different primer pairs: SIV-ampA-F (5’ TGTCTTTTATCCAGGAAGGGGTA 3’), SIV-ampA-R (5’ CTCTAATTAACCTACAGAGATGTTTGG 3’), SIV-ampB-F (5’ AAAATTGAAGCAGTGGCCATTAT 3’), SIV-ampB-R (5’ TACTTATGAGCTCTCGGGAACCT 3’), SIV-ampC-F (5’ GCTTTACAGCGGGAGAAGTG 3’), SIV-ampC-R (5’ TGCCAAGTGTTGATTATTTGTC 3’), SIV-ampD-F (5’ GGTGTTGGTTTGGAGGAAAA 3’), SIV-ampD-R (5’ GAATACAGAGCGAAATGCAGTG 3’). The four amplicons were purified and then quantified using a Quant-IT double-stranded DNA (dsDNA) HS kit (Invitrogen). The four amplicons were pooled to a total of 1ng and then the Nextera XT kit (Illumina) was used to fragment the DNA and generate uniquely tagged libraries. The tagged libraries were quantified and fragment size distribution was determined with a high-sensitivity Agilent Bioanalyzer chip. Libraries were pooled and sequenced on an Illumina MiSeq. The sequence confirmation of the SIV_mac_239 K180S challenge stock and *env* sequences in plasma collected at weeks 4 and 8 post-infection in macaques challenged with this virus are available at SRA BioProject PRJNA1031618.

### Statistical analysis

Macaques were randomly assigned to treatment or control groups that received the Env-specific antibody PGT145 or the control antibody DEN3. In two independent experiments, the animals were challenged intrarectally with SIVmac239 or SIVmac239 K180S. Statistical analyses were performed separately for each challenge experiment. Viral loads and CD4^+^ T cell counts were log10 transformed prior to statistical modelling to create more normally distributed datasets. Mean log10 peak viral loads and time to peak viremia were compared between antibody treatment and control groups by linear regression. Differences in chronic phase viral loads (8-24 weeks) between antibody treatment and control groups were compared using linear mixed models adjusted for repeated measurements on an animal.

## Acknowledgments

We thank the Quantitative Molecular Diagnostics Core of the AIDS and Cancer Virus Program, Frederick National Laboratory, for expert assistance with plasma viral load determinations. D.T.E is an Elizabeth Glaser Scientist of the Elizabeth Glaser Pediatric AIDS Foundation (www.pedaids.org).

## Supporting information

**S1 Fig. Resistance of the SIVmac239 K180S challenge virus to neutralization by PGT145.** Two-fold dilutions of the SIVmac239 K180S challenge virus (starting at 5.56 ng/ml p27) were incubated in the presence of a constant amount of PGT145 and K11 (50 µg/ml) for one hour before the addition of TZM-bl cells. Luciferase activity in the TZM-bl cells as an indicator of SIV infectivity was measured after a 3-day incubation. The percentage of residual infectivity was calculated from the luciferase activity in presence of each antibody relative to maximal luciferase activity in the absence of antibody after subtracting background luciferase in uninfected cells.

**S2 Fig. Serum concentrations of PGT145 decline after passive administration.** Serum concentrations of PGT145 in animals challenged with SIV_mac_239 K180S were measured by ELISA on plates coated with an anti-His antibody and captured 6-His-tagged HIV-1 BG505 SOSIP trimers. Error bars indicate standard deviation of the mean. The data was analyzed by nonlinear regression. The best-fit cure is shown in black with an R^2^-value of 0.9698 and an estimate antibody half-life of 8.7 days.

## References

1. Rerks-Ngarm S, Pitisuttithum P, Nitayaphan S, Kaewkungwal J, Chiu J, Paris R, et al. Vaccination with ALVAC and AIDSVAX to prevent HIV-1 infection in Thailand. N Engl J Med. 2009;361(23):2209–20. doi: 10.1056/NEJMoa0908492. PubMed PMID: 19843557.

2. Haynes BF, Gilbert PB, McElrath MJ, Zolla-Pazner S, Tomaras GD, Alam SM, et al. Immune-correlates analysis of an HIV-1 vaccine efficacy trial. N Engl J Med. 2012;366:1275–86.

3. Rolland M, Edlefsen PT, Larsen BB, Tovanabutra S, Sanders-Buell E, Hertz T, et al. Increased HIV-1 vaccine efficacy against viruses with genetic signatures in Env V2. Nature. 2012;490(7420):417-20. Epub 20120910. doi: 10.1038/nature11519. PubMed PMID: 22960785; PubMed Central PMCID: PMCPMC3551291.

4. Yates NL, Liao HX, Fong Y, deCamp A, Vandergrift NA, Williams WT, et al. Vaccine-induced Env V1-V2 IgG3 correlates with lower HIV-1 infection risk and declines soon after vaccination. Sci Transl Med. 2014;6(228):228ra39. doi: 10.1126/scitranslmed.3007730. PubMed PMID: 24648342; PubMed Central PMCID: PMC4116665.

5. Chung AW, Ghebremichael M, Robinson H, Brown E, Choi I, Lane S, et al. Polyfunctional Fc-effector profiles mediated by IgG subclass selection distinguish RV144 and VAX003 vaccines. Sci Transl Med. 2014;6(228):228ra38. doi: 10.1126/scitranslmed.3007736. PubMed PMID: 24648341.

6. Desrosiers RC. Protection against HIV Acquisition in the RV144 Trial. J Virol. 2017;91(18). Epub 20170824. doi: 10.1128/JVI.00905-17. PubMed PMID: 28701398; PubMed Central PMCID: PMCPMC5571267.

7. Gray GE, Bekker LG, Laher F, Malahleha M, Allen M, Moodie Z, et al. Vaccine Efficacy of ALVAC-HIV and Bivalent Subtype C gp120-MF59 in Adults. N Engl J Med. 2021;384(12):1089–100. doi: 10.1056/NEJMoa2031499. PubMed PMID: 33761206; PubMed Central PMCID: PMCPMC7888373.

8. Barouch DH, Liu J, Li H, Maxfield LF, Abbink P, Lynch DM, et al. Vaccine protection against acquisition of neutralization-resistant SIV challenges in rhesus monkeys. Nature. 2012;482(7383):89-93. doi: 10.1038/nature10766. PubMed PMID: 22217938; PubMed Central PMCID: PMC3271177.

9. Gordon SN, Doster MN, Kines RC, Keele BF, Brocca-Cofano E, Guan Y, et al. Antibody to the gp120 V1/V2 loops and CD4+ and CD8+ T cell responses in protection from SIVmac251 vaginal acquisition and persistent viremia. J Immunol. 2014;193(12):6172–83. Epub 20141114. doi: 10.4049/jimmunol.1401504. PubMed PMID: 25398324; PubMed Central PMCID: PMCPMC4335709.

10. Gordon SN, Liyanage NPM, Doster MN, Vaccari M, Vargas-Inchaustegui DA, Pegu P, et al. Boosting of ALVAC-SIV Vaccine-Primed Macaques with the CD4-SIVgp120 Fusion Protein Elicits Antibodies to V2 Associated with a Decreased Risk of SIVmac251 Acquisition. J Immunol. 2016;197(7):2726–37. doi: 10.4049/jimmunol.1600674. PubMed PMID: WOS:000385004600022.

11. Vaccari M, Gordon SN, Fourati S, Schifanella L, Liyanage NP, Cameron M, et al. Adjuvant-dependent innate and adaptive immune signatures of risk of SIVmac251 acquisition. Nat Med. 2016;22(7):762–70. Epub 2016/05/31. doi: 10.1038/nm.4105. PubMed PMID: 27239761; PubMed Central PMCID: PMCPMC5916782.

12. Vaccari M, Fourati S, Gordon SN, Brown DR, Bissa M, Schifanella L, et al. HIV vaccine candidate activation of hypoxia and the inflammasome in CD14(+) monocytes is associated with a decreased risk of SIVmac251 acquisition. Nat Med. 2018;24(6):847–56. Epub 2018/05/23. doi: 10.1038/s41591-018-0025-7. PubMed PMID: 29785023; PubMed Central PMCID: PMCPMC5992093.

13. Silva de Castro I, Gorini G, Mason R, Gorman J, Bissa M, Rahman MA, et al. Anti-V2 antibodies virus vulnerability revealed by envelope V1 deletion in HIV vaccine candidates. iScience. 2021;24(2):102047. Epub 20210109. doi: 10.1016/j.isci.2021.102047. PubMed PMID: 33554060; PubMed Central PMCID: PMCPMC7847973.

14. Barouch DH, Stephenson KE, Borducchi EN, Smith K, Stanley K, McNally AG, et al. Protective efficacy of a global HIV-1 mosaic vaccine against heterologous SHIV challenges in rhesus monkeys. Cell. 2013;155(3):531–9. Epub 20131024. doi: 10.1016/j.cell.2013.09.061. PubMed PMID: 24243013; PubMed Central PMCID: PMCPMC3846288.

15. Pegu P, Vaccari M, Gordon S, Keele BF, Doster M, Guan Y, et al. Antibodies with high avidity to the gp120 envelope protein in protection from simian immunodeficiency virus SIV(mac251) acquisition in an immunization regimen that mimics the RV-144 Thai trial. J Virol. 2013;87(3):1708–19. Epub 20121121. doi: 10.1128/JVI.02544-12. PubMed PMID: 23175374; PubMed Central PMCID: PMCPMC3554145.

16. Barouch DH, Alter G, Broge T, Linde C, Ackerman ME, Brown EP, et al. Protective efficacy of adenovirus/protein vaccines against SIV challenges in rhesus monkeys. Science. 2015;349(6245):320-4. Epub 20150702. doi: 10.1126/science.aab3886. PubMed PMID: 26138104; PubMed Central PMCID: PMCPMC4653134.

17. Bradley T, Pollara J, Santra S, Vandergrift N, Pittala S, Bailey-Kellogg C, et al. Pentavalent HIV-1 vaccine protects against simian-human immunodeficiency virus challenge. Nature communications. 2017;8:15711. Epub 2017/06/09. doi: 10.1038/ncomms15711. PubMed PMID: 28593989; PubMed Central PMCID: PMCPMC5472724.

18. Hessell AJ, Li L, Malherbe DC, Barnette P, Pandey S, Sutton W, et al. Virus Control in Vaccinated Rhesus Macaques Is Associated with Neutralizing and Capturing Antibodies against the SHIV Challenge Virus but Not with V1V2 Vaccine-Induced Anti-V2 Antibodies Alone. J Immunol. 2021;206(6):1266–83. Epub 20210203. doi: 10.4049/jimmunol.2001010. PubMed PMID: 33536254; PubMed Central PMCID: PMCPMC7946713.

19. Hessell AJ, Shapiro MB, Powell R, Malherbe DC, McBurney SP, Pandey S, et al. Reduced Cell-Associated DNA and Improved Viral Control in Macaques following Passive Transfer of a Single Anti-V2 Monoclonal Antibody and Repeated Simian/Human Immunodeficiency Virus Challenges. J Virol. 2018;92(11). Epub 20180514. doi: 10.1128/JVI.02198-17. PubMed PMID: 29514914; PubMed Central PMCID: PMCPMC5952134.

20. Alter G, Yu WH, Chandrashekar A, Borducchi EN, Ghneim K, Sharma A, et al. Passive Transfer of Vaccine-Elicited Antibodies Protects against SIV in Rhesus Macaques. Cell. 2020;183(1):185–96 e14. Epub 2020/10/03. doi: 10.1016/j.cell.2020.08.033. PubMed PMID: 33007262; PubMed Central PMCID: PMCPMC7534693.

21. Fuchs SP, Martinez-Navio JM, Piatak M, Jr., Lifson JD, Gao G, Desrosiers RC. AAV-Delivered Antibody Mediates Significant Protective Effects against SIVmac239 Challenge in the Absence of Neutralizing Activity. PLoS Pathog. 2015;11(8):e1005090. Epub 2015/08/08. doi: 10.1371/journal.ppat.1005090. PubMed PMID: 26248318; PubMed Central PMCID: PMCPMC4527674.

22. Astronomo RD, Santra S, Ballweber-Fleming L, Westerberg KG, Mach L, Hensley-McBain T, et al. Neutralization Takes Precedence Over IgG or IgA Isotype-related Functions in Mucosal HIV-1 Antibody-mediated Protection. EBioMedicine. 2016;14:97–111. doi: 10.1016/j.ebiom.2016.11.024. PubMed PMID: 27919754; PubMed Central PMCID: PMCPMC5161443.

23. Burton DR, Hessell AJ, Keele BF, Klasse PJ, Ketas TA, Moldt B, et al. Limited or no protection by weakly or nonneutralizing antibodies against vaginal SHIV challenge of macaques compared with a strongly neutralizing antibody. Proc Natl Acad Sci U S A. 2011;108(27):11181–6. Epub 20110620. doi: 10.1073/pnas.1103012108. PubMed PMID: 21690411; PubMed Central PMCID: PMCPMC3131343.

24. Santra S, Tomaras GD, Warrier R, Nicely NI, Liao HX, Pollara J, et al. Human Non-neutralizing HIV-1 Envelope Monoclonal Antibodies Limit the Number of Founder Viruses during SHIV Mucosal Infection in Rhesus Macaques. PLoS Pathog. 2015;11(8):e1005042. doi: 10.1371/journal.ppat.1005042. PubMed PMID: 26237403; PubMed Central PMCID: PMCPMC4523205.

25. von Bredow B, Arias JF, Heyer LN, Moldt B, Le K, Robinson JE, et al. Comparison of antibody-dependent cell-mediated cytotoxicity and virus neutralization by HIV-1 Env-specific monoclonal antibodies. J Virol. 2016;90(13):6127–39. doi: 10.1128/JVI.00347-16. PubMed PMID: 27122574; PubMed Central PMCID: PMCPMC4907221.

26. Bruel T, Guivel-Benhassine F, Amraoui S, Malbec M, Richard L, Bourdic K, et al. Elimination of HIV-1-infected cells by broadly neutralizing antibodies. Nature Communication. 2016;7:10844. doi: 10.1038/ncomms10844. PubMed PMID: 26936020; PubMed Central PMCID: PMCPMC4782064.

27. Bruel T, Guivel-Benhassine F, Lorin V, Lortat-Jacob H, Baleux F, Bourdic K, et al. Lack of ADCC Breadth of Human Nonneutralizing Anti-HIV-1 Antibodies. J Virol. 2017;91(8). doi: 10.1128/JVI.02440-16. PubMed PMID: 28122982; PubMed Central PMCID: PMCPMC5375671.

28. Ren Y, Korom M, Truong R, Chan D, Huang SH, Kovacs CC, et al. Susceptibility to Neutralization by Broadly Neutralizing Antibodies Generally Correlates with Infected Cell Binding for a Panel of Clade B HIV Reactivated from Latent Reservoirs. J Virol. 2018;92(23). Epub 2018/09/14. doi: 10.1128/JVI.00895-18. PubMed PMID: 30209173; PubMed Central PMCID: PMCPMC6232479.

29. Prevost J, Anand SP, Rajashekar JK, Zhu L, Richard J, Goyette G, et al. HIV-1 Vpu restricts Fc-mediated effector functions in vivo. Cell Rep. 2022;41(6):111624. doi: 10.1016/j.celrep.2022.111624. PubMed PMID: 36351384; PubMed Central PMCID: PMCPMC9703018.

30. Grunst MW, Ladd RA, Clark NM, Gil HM, Klenchin VA, Mason R, et al. Antibody-dependent cellular cytotoxicity, infected cell binding and neutralization by antibodies to the SIV envelope glycoprotein. PLoS Pathog. 2023;19(5):e1011407. Epub 20230530. doi: 10.1371/journal.ppat.1011407. PubMed PMID: 37253062; PubMed Central PMCID: PMCPMC10256149.

31. Lee JH, Andrabi R, Su CY, Yasmeen A, Julien JP, Kong L, et al. A broadly neutralizing antibody targets the dynamic HIV envelope trimer apex via a long, rigidified, and anionic beta-hairpin structure. Immunity. 2017;46(4):690–702. doi: 10.1016/j.immuni.2017.03.017. PubMed PMID: 28423342; PubMed Central PMCID: PMCPMC5400778.

32. von Bredow B, Andrabi R, Grunst M, Grandea AG, 3rd, Le K, Song G, et al. Differences in the Binding Affinity of an HIV-1 V2 Apex-Specific Antibody for the SIVsmm/mac Envelope Glycoprotein Uncouple Antibody-Dependent Cellular Cytotoxicity from Neutralization. MBio. 2019;10(4). Epub 2019/07/04. doi: 10.1128/mBio.01255-19. PubMed PMID: 31266872; PubMed Central PMCID: PMCPMC6606807.

33. Hoffenberg S, Powell R, Carpov A, Wagner D, Wilson A, Kosakovsky Pond S, et al. Identification of an HIV-1 clade A envelope that exhibits broad antigenicity and neutralization sensitivity and elicits antibodies targeting three distinct epitopes. J Virol. 2013;87(10):5372–83. Epub 20130306. doi: 10.1128/JVI.02827-12. PubMed PMID: 23468492; PubMed Central PMCID: PMCPMC3648150.

34. Alpert MD, Heyer LN, Williams DEJ, Harvey JD, Greenough T, Allhorn M, et al. A novel assay for antibody-dependent cell-mediated cytotoxicity against HIV-1- or SIV-infected cells reveals incomplete overlap with antibodies measured by neutralization and binding assays. J Virol. 2012;86:12039–52.

35. Carias AM, Schneider JR, Madden P, Lorenzo-Redondo R, Arainga M, Pegu A, et al. Anatomic Distribution of Intravenously Injected IgG Takes Approximately 1 Week to Achieve Stratum Corneum Saturation in Vaginal Tissues. J Immunol. 2021;207(2):505–11. Epub 20210623. doi: 10.4049/jimmunol.2100253. PubMed PMID: 34162723; PubMed Central PMCID: PMCPMC8516693.

36. Pegu A, Borate B, Huang Y, Pauthner MG, Hessell AJ, Julg B, et al. A Meta-analysis of Passive Immunization Studies Shows that Serum-Neutralizing Antibody Titer Associates with Protection against SHIV Challenge. Cell Host Microbe. 2019;26(3):336–46 e3. Epub 2019/09/13. doi: 10.1016/j.chom.2019.08.014. PubMed PMID: 31513771; PubMed Central PMCID: PMCPMC6755677.

37. Zhao F, Berndsen ZT, Pedreno-Lopez N, Burns A, Allen JD, Barman S, et al. Molecular insights into antibody-mediated protection against the prototypic simian immunodeficiency virus. Nature communications. 2022;13(1):5236. Epub 20220906. doi: 10.1038/s41467-022-32783-2. PubMed PMID: 36068229; PubMed Central PMCID: PMCPMC9446601.

38. McCoy LE, Falkowska E, Doores KJ, Le K, Sok D, van Gils MJ, et al. Incomplete Neutralization and Deviation from Sigmoidal Neutralization Curves for HIV Broadly Neutralizing Monoclonal Antibodies. PLoS Pathog. 2015;11(8):e1005110. Epub 20150812. doi: 10.1371/journal.ppat.1005110. PubMed PMID: 26267277; PubMed Central PMCID: PMCPMC4534392.

39. Doores KJ, Burton DR. Variable loop glycan dependency of the broad and potent HIV-1-neutralizing antibodies PG9 and PG16. J Virol. 2010;84(20):10510–21. Epub 20100804. doi: 10.1128/JVI.00552-10. PubMed PMID: 20686044; PubMed Central PMCID: PMCPMC2950566.

40. Gorman J, Wang C, Mason RD, Nazzari AF, Welles HC, Zhou T, et al. Cryo-EM structures of prefusion SIV envelope trimer. Nat Struct Mol Biol. 2022;29(11):1080–91. Epub 20221107. doi: 10.1038/s41594-022-00852-1. PubMed PMID: 36344847.

41. Valley CC, Cembran A, Perlmutter JD, Lewis AK, Labello NP, Gao J, et al. The methionine-aromatic motif plays a unique role in stabilizing protein structure. J Biol Chem. 2012;287(42):34979–91. Epub 20120801. doi: 10.1074/jbc.M112.374504. PubMed PMID: 22859300; PubMed Central PMCID: PMCPMC3471747.

42. Corey L, Gilbert PB, Juraska M, Montefiori DC, Morris L, Karuna ST, et al. Two Randomized Trials of Neutralizing Antibodies to Prevent HIV-1 Acquisition. N Engl J Med. 2021;384(11):1003–14. doi: 10.1056/NEJMoa2031738. PubMed PMID: 33730454; PubMed Central PMCID: PMCPMC8189692.

43. Shingai M, Donau OK, Plishka RJ, Buckler-White A, Mascola JR, Nabel GJ, et al. Passive transfer of modest titers of potent and broadly neutralizing anti-HIV monoclonal antibodies block SHIV infection in macaques. J Exp Med. 2014;211(10):2061–74. doi: 10.1084/jem.20132494. PubMed PMID: 25155019; PubMed Central PMCID: PMC4172223.

44. Garber JC, Barbee RW, Bielitzki JT, Clayton LA, Donovan JC, Hendricksen CFM, et al. Guide for the care and use of laboratory animals. 8th Edition ed. Washington, DC: The National Academies Press; 2011.

45. Kozlowski PA, Lynch RM, Patterson RR, Cu-Uvin S, Flanigan TP, Neutra MR. Modified wick method using Weck-Cel sponges for collection of human rectal secretions and analysis of mucosal HIV antibody. J Acquir Immune Defic Syndr. 2000;24:297–309.

46. Wei X, Decker JM, Liu H, Zhang Z, Arani RB, Kilby JM, et al. Emergence of resistant human immunodeficiency virus type 1 in patients receiving fusion inhibitor (T-20) monotherapy. Antimicrob Agents Chemother. 2002;46(6):1896–905. PubMed PMID: 12019106; PubMed Central PMCID: PMC127242.

47. Wei X, Decker JM, Wang S, Hui H, Kappes JC, Wu X, et al. Antibody neutralization and escape by HIV-1. Nature. 2003;422(6929):307-12. doi: 10.1038/nature01470. PubMed PMID: 12646921.

48. Alpert MD, Harvey JD, Lauer WA, Reeves RK, Michael Piatak J, Carville A, et al. ADCC develops over time during peristent infection with live-attenuated SIV and is associate with complete protection against SIVmac251 challenge. PLoS Pathog. 2012;8:e1002890.

49. Li H, Wang S, Kong R, Ding W, Lee FH, Parker Z, et al. Envelope residue 375 substitutions in simian-human immunodeficiency viruses enhance CD4 binding and replication in rhesus macaques. Proc Natl Acad Sci U S A. 2016;113(24):E3413–22. Epub 2016/06/02. doi: 10.1073/pnas.1606636113. PubMed PMID: 27247400; PubMed Central PMCID: PMCPMC4914158.

50. Sutton MS, Ellis-Connell A, Moriarty RV, Balgeman AJ, Gellerup D, Barry G, et al. Acute-Phase CD4(+) T Cell Responses Targeting Invariant Viral Regions Are Associated with Control of Live Attenuated Simian Immunodeficiency Virus. J Virol. 2018;92(21). Epub 20181012. doi: 10.1128/JVI.00830-18. PubMed PMID: 30111562; PubMed Central PMCID: PMCPMC6189504.

51. Goddard TD, Huang CC, Meng EC, Pettersen EF, Couch GS, Morris JH, et al. UCSF ChimeraX: Meeting modern challenges in visualization and analysis. Protein Sci. 2018;27(1):14–25. Epub 20170906. doi: 10.1002/pro.3235. PubMed PMID: 28710774; PubMed Central PMCID: PMCPMC5734306.

